# Epigenetic Age-Predictor for Mice based on Three CpG Sites

**DOI:** 10.1101/296525

**Authors:** Yang Han, Monika Eipel, Vadim Sakk, Bertien Dethmers-Ausema, Gerald de Haan, Hartmut Geiger, Wolfgang Wagner

## Abstract

Epigenetic clocks for mice were generated based on deep-sequencing analysis of the methylome. In this study we demonstrate that site-specific analysis of DNA methylation levels by pyrosequencing at only three CG dinucleotides (CpGs) in the genes *Prima1*, *Hsf4*, and *Kcns1* facilitates precise estimation of chronological age in murine blood samples, too. DBA/2J mice revealed accelerated epigenetic aging as compared to C57BL6 mice, which is in line with their shorter life-expectancy. The three-CpG-predictor provides a simple and cost-effective biomarker to determine biological age in large intervention studies with mice.

## Introduction

Age-associated DNA methylation (DNAm) was first described for humans after Illumina Bead Chip microarray data became available to enable cross comparison of thousands of CpG loci (Bocklandt *et al.* 2011; Koch & Wagner 2011). Many of these age-associated CpGs were then integrated into epigenetic age-predictors (Hannum *et al.* 2013; Horvath 2013; Weidner *et al.* 2014). However, site-specific DNAm analysis at individual CpGs can also provide robust biomarkers for aging. For example, we have described that DNAm analysis at only three CpGs enables age-predictions for human blood samples with a mean absolute deviation (MAD) from chronological age of less than five years (Weidner *et al.* 2014). Such simplistic age-predictors for human specimen are widely used because they enable fast and cost-effective analysis in large cohorts.

Recently, epigenetic clocks were also published for mice by using either reduced representation bisulfite sequencing (RRBS) or whole genome bisulfite sequencing (WGBS) (Petkovich *et al.* 2017; Stubbs *et al.* 2017; Wang *et al.* 2017). For example, Petkovich et al. described a 90 CpG model for blood (Petkovich *et al.* 2017), and Stubbs and coworkers a 329 CpG model for various different tissues (Stubbs *et al.* 2017). Nutrition and genetic background seem to affect the epigenetic age of mice – and thereby possibly aging of the organism (Cole *et al.* 2017; Hahn *et al.* 2017; Maegawa *et al.* 2017). In analogy, epigenetic aging of humans is associated with life expectancy, indicating that it rather reflects biological age than chronological age (Marioni *et al.* 2015; Lin *et al.* 2016). However, DNAm profiling by deep sequencing technology is technically still challenging, relatively expensive, and not every sequencing-run covers all relevant CpG sites with enough reading depth.

## Results and Discussion

Therefore, we established pyrosequencing assays for nine genomic regions of previously published predictors (Petkovich *et al.* 2017; Stubbs *et al.* 2017). These regions were preselected to have multiple age-associated CpGs in close vicinity. DNAm was then analyzed in 24 blood samples of female C57BL/6 mice (11 to 117 weeks old; Table S1). Three CpGs revealed high correlation with chronological age, and were associated with the genes Proline rich membrane anchor 1 (*Prima1:* chr12:103214639; R^2^ = 0.71), Heat shock transcription factor 4 (*Hsf4:* chr8:105271000; R^2^ = 0.95) and Potassium voltage-gated channel modifier subfamily S member 1 (*Kcns1:* chr2:164168110; R^2^ = 0.83; Figure 1A-C). Notably, all three CpGs were derived from the epigenetic age-predictor for blood samples (Petkovich *et al.* 2017). A multivariate model for age-predictions was established for DNAm at the CpGs in *Prima 1* (α), *Hsf4* (β), and *Kcns1* (γ):

**Figure 1.**
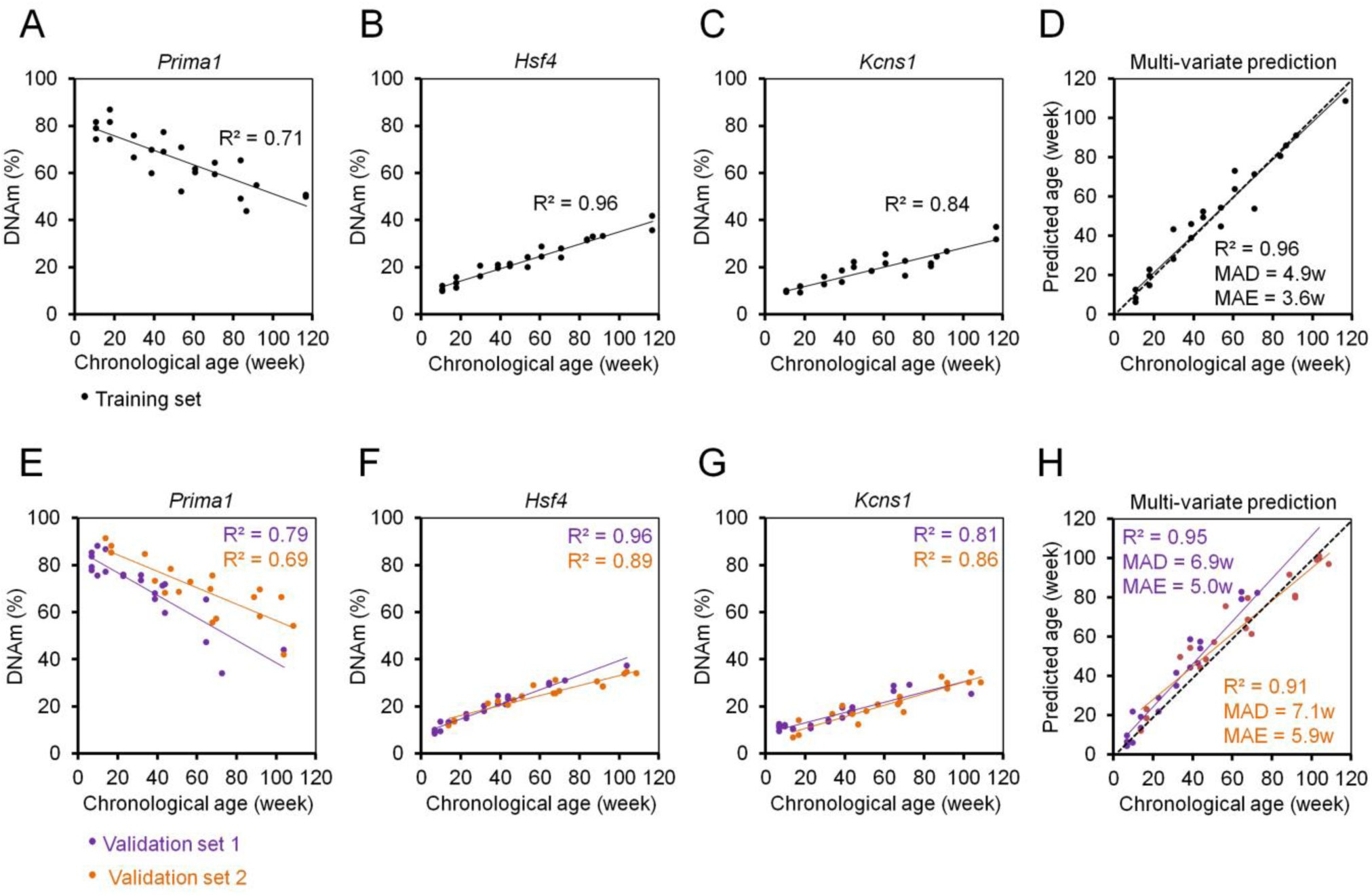
Three CpG epigenetic age-predictor for mice. **(A-C)** DNA methylation (DNAm) of three CpGs in the genes *Prima1*, *Hsf4* and *Kcns1* was analyzed by pyrosequening in 24 C57BL/6 mice (training set). Spearman correlation (R^2^) of DNAm *versus* chronological age is indicated. **(D)** Based on these age-associated DNAm changes a multivariate model for age prediction was calculated. **(E-G)** Subsequently, two independent validation sets were analyzed: 21 C57BL/6 mice from the University of Ulm and 19 C57BL/6 mice from the University of Groningen (validation sets 1 and 2, respectively). **(H)** Age predictions with the three-CpG-model revealed a high correlation with chronological age in the independent validation sets (MAD = mean absolute deviation; MAE = median absolute error).

Predicted age^C57BL/6^ (in weeks) = -58.076 + 0.25788 α + 3.06845 β + 1.00879 γ

Age-predictions correlated very well with the chronological age of C57BL/6 mice in the training set (R^2^ = 0.96; MAD = 4.86 weeks; Figure 1D) and this was subsequently validated in a blinded manner for 21 C57BL/6J mice (7 to 104 weeks old) from the University of Ulm (validation set 1) and 19 C57BL/6J mice (14 to 109 weeks old) from the University of Groningen (validation set 2). The results of both validation sets revealed high correlations with chronological age (R^2^ = 0.95 and 0.91, respectively; Figure 1E-H) with relatively small MADs (6.9 and 7.1 weeks) and median absolute errors (MAE; 5.0 and 5.9 weeks). Thus, our age-predictions seem to have similar precision as previously described for multi-CpG predictors based on RRBS or WGBS data (Petkovich *et al.* 2017; Stubbs *et al.* 2017; Wang *et al.* 2017).

Gender did not have significant impact on our epigenetic age-predictions for mice (Figure S1), as described before (Maegawa *et al.* 2017; Petkovich *et al.* 2017; Stubbs *et al.* 2017). In contrast, the human epigenetic clock is clearly accelerated in male donors (Hannum *et al.* 2013; Horvath 2013; Weidner *et al.* 2014). This coincides with shorter life expectancy in men than woman, whereas in mice there are no consistent sex differences in longevity (Goodrick 1975).

Subsequently, we analyzed epigenetic aging of DBA/2 mice that have a shorter life expectancy than C57BL/6 mice (Goodrick 1975) (25 mice from Ulm and Groningen; 7 to 109 weeks old). The three CpGs in *Prima1*, *Hsf4* and *Kcns1* revealed high correlation with chronological age (R^2^= 0.83, 0.76 and 0.64, respectively), albeit the different slopes indicated accelerated epigenetic age for each of the CpGs (Figure S2A-C). Conversely, epigenetic age-predictions were significantly overestimated in the shorter-lived DBA/2 mice (*P* < 0.0001; Figure 2). These results provide further evidence that age-associated DNAm is generally rather related to biological age and that age-predictors should be adjusted for different inbreed mice strains. To this end, we have retrained a multivariate model for DBA/2 mice:

**Figure 2.**
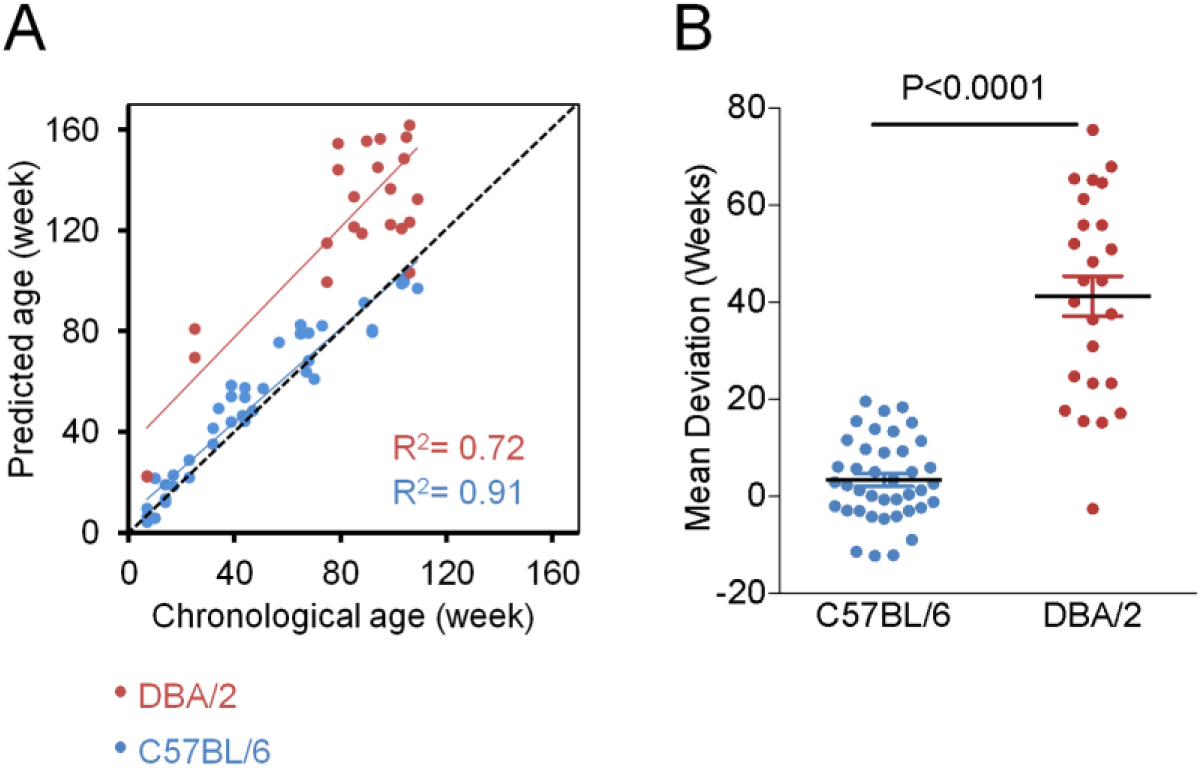
Epigenetic aging is accelerated in DBA/2 mice as compared to C57BL/6 mice. **(A)** Epigenetic age-predictions based on three CpGs for blood samples of C57BL/6 (blue) and DBA/2 mice (red; multivariate model for C57BL/6; R^2^ = Spearman correlation). **(B)** Mean deviations (MAD) demonstrate that the epigenetic age is significantly overestimated in DBA/2 mice (Mann–Whitney U test).

Predicted age^DBA/2^ (in weeks) = 91.04858 - 1.21037 α + 1.02067 β + 0.22891 γ

This adjusted model facilitated relatively precise age-predictions for DBA/2 mice (R^2^ = 0.89; MAD = 8.5 weeks; MAE = 8.3 weeks; Figure S2D). However, in elderly DBA/2 mice the epigenetic age predictions revealed higher “errors” from chronological age, which might be attributed to the fact that the variation of lifespan is higher in DBA/2 than C57BL/6 mice (Goodrick 1975; de Haan *et al.* 1998).

Taken together, we describe an easily applicable but quite precise approach to determine epigenetic age of mice. We believe that our assay will be instrumental to gain additional insight into mechanisms that regulate age-associated DNAm and for longevity intervention studies in mice.

## Experimental Procedures

### Mouse strains and blood collection

Blood samples of C57BL/6J mice of the training set and of the validation set 1 were taken at the University of Ulm by submandibular bleeding (100-200 μl) of living mice or postmortem from the vena cava. C57BL/6J samples of the validation set 2 were taken at the University of Groningen from the cheek. DBA/2J samples were taken at the University of Ulm (n = 6) and Groningen (n = 19). All mice were accommodated under pathogen-free conditions. Experiments were approved by the Institutional Animal Care of the Ulm University as well as by Regierungspräsidium Tübingen and by the Institutional Animal Care and Use Committee of the University of Groningen (IACUC-RUG), respectively.

### Genomic DNA isolation and bisulfite conversion

Genomic DNA was isolated from 50 µl blood using the QIAamp DNA Mini Kit (Qiagen, Hilden, Germany). DNA concentration was quantified by Nanodrop 2000 Spectrophotometers (Thermo Scientific, Wilmington, USA). 200 ng of genomic DNA was subsequently bisulfite-converted with the EZ DNA Methylation Kit (Zymo Research, Irvine, USA).

### Pyrosequencing

Bisulfite converted DNA was subjected to PCR amplification. Primers were purchased at Metabion and the sequences are provided in Table S2. 20 µg PCR products was immobilized to 5 µl Streptavidin Sepharose High Performance Bead (GE Healthcare, Piscataway, NJ, USA), and then was annealed to 1 µl sequencing primer (5 μM) for 2 minutes at 80°C. Amplicons were sequenced on PyroMark Q96 ID System (Qiagen, Hilden, Germany) and analyzed with PyroMark Q CpG software (Qiagen). The relevant sequences are depicted for the three relevant genomic regions in Figure S3.

### Statistical analysis

Linear regressions, MAD and MAE were calculated with Excel. Statistical significance of the deviations between predicted and chronological age was estimated by Mann–Whitney U test.

## Supplemental information

The supplemental PDF comprises Figures S1-S3 and Tables S1-S2.

## Authors contributions

Y.H. and M.E. performed pyrosequencing and analyzed the data; V.S., B.D.-A., G.d.H., and H.G. provided samples and supported the study design; W.W. conceived the study and analyzed the data, Y.H. and W.W. wrote the manuscript and all authors read and approved the final manuscript.

